# Developmental Trajectory of Synaptic Remodeling in the Mouse Prefrontal Cortex

**DOI:** 10.64898/2026.04.26.720930

**Authors:** Johanna Furrer, Sarai Fischer, Alexandra von Faber-Castell, Sina M. Schalbetter, Viktor Beilmann, Eliane Boesch, Matthias T. Wyss, Urs Meyer, Bruno Weber, Tina Notter

## Abstract

The prefrontal cortex (PFC) is a heteromodal association area critical for higher-order cognitive functions. Its protracted maturation extends through adolescence into early adulthood, a period characterized by extensive remodeling of neuronal networks and synaptic architecture. This heightened plasticity supports the refinement of prefrontal circuits necessary to meet the evolving cognitive and behavioral demands of this developmental transition. However, the extended maturation window also increases vulnerability to environmental perturbations, which can lead to lasting synaptic and cognitive impairments. Despite widespread use of developmental disruption models during adolescence, a systematic characterization of synaptic dynamics during normal PFC maturation is lacking. Here, we combined longitudinal *in vivo* two-photon imaging of dendritic spines with cross-sectional quantification of excitatory and inhibitory synapses, and microglia-mediated synaptic engulfment, in mice from juvenile to adult stages. This integrated approach provides a comprehensive reference atlas of normal prefrontal synaptic maturation, offering a framework for interpreting alterations to synapses in models of developmental disturbance.

## INTRODUCTION

The prefrontal cortex (PFC) is a cortical association area that subserves many fundamental higher-order cognitive functions, including decision-making, goal-directed behavior, cognitive flexibility, attention, and working memory(Carlén, 2017; Friedman and Robbins, 2022). A defining feature of the PFC is its protracted maturation, which extends throughout adolescence into early adulthood, making it the last brain region to reach full structural and functional maturity. This prolonged developmental trajectory is evolutionarily conserved and has been documented across species, including humans(Pfefferbaum et al., 1994; Gogtay et al., 2004; O’Donnell et al., 2005), non-human primates(Xia et al., 2023), and rodents(Juraska and Drzewiecki, 2020).

During adolescence, the PFC undergoes extensive remodeling of its neuronal networks and synaptic architecture(Juraska and Drzewiecki, 2020; Chini and Hanganu-Opatz, 2021). This period of heightened synaptic plasticity is thought to support the refinement of prefrontal circuits, enabling adaptive responses to the evolving cognitive and behavioral demands associated with the transition from adolescence to adulthood(Paus, 2005; Larsen and Luna, 2018; Klune et al., 2021). However, this extended developmental window also confers increased vulnerability to perturbations(Feinberg, 1982; Andersen, 2003). Consistent with this notion, experimental studies in rodents suggest that environmental adversities during this critical period can lead to persistent synaptic and cognitive impairments in adulthood(Hinton et al., 2019; Miller et al., 2019; Shaw et al., 2020; Benoit et al., 2022; Schalbetter et al., 2022; Pöpplau et al., 2024).

Despite widespread use of developmental disruption models in preclinical psychiatric research, a systematic characterization of normal postnatal synaptic maturation is still missing. Work in humans(Huttenlocher, 1979; Petanjek et al., 2011), non-human primates(Bourgeois et al., 1994; Anderson et al., 1995; Elston et al., 2009), and rodents(Gourley et al., 2012; Koss et al., 2014; Drzewiecki et al., 2016; Delevich et al., 2020; Pöpplau et al., 2024) demonstrates that prefrontal synaptic density declines from adolescence to adulthood, but these studies usually measure only presynaptic terminals or postsynaptic spines and rarely distinguish excitatory from inhibitory synapses. Most previous studies have relied on cross-sectional, post-mortem analyses, lacking critical insight into the dynamics of synapse formation and elimination within the same individual. As a result, a comprehensive, longitudinal profile of normative prefrontal synaptic maturation remains missing, limiting interpretation of experiments targeting synaptic remodeling from juvenile to adult stages.

To address this limitation, we systematically quantified synaptic remodeling in the mouse PFC across juvenile to adult stages. We combined longitudinal, high-resolution *in vivo* two-photon imaging to track dendritic spine dynamics within the same individuals with cross-sectional quantification of excitatory and inhibitory synapse densities across matching ages. This integrated approach yields a comprehensive profile of normative prefrontal synaptic maturation in male and female mice, establishing a reference framework for interpreting synaptic perturbations in developmental disruption models of psychiatric disorders.

## RESULTS

### Developmental trajectory of *in vivo* spine dynamics across prefrontal maturation

We performed high-resolution, longitudinal *in vivo* two-photon imaging to track spine dynamics in the maturing PFC from early adolescence to adulthood. Thy1-GFP-M mice for sparse neuronal labeling(Feng et al., 2000) were used to visualize individual dendrites and spines over time. To access medial PFC (mPFC), 3-week-old Thy1-GFP-M mice were implanted with a right-angled microprism attached to a cranial window into the subdural space of the contralateral hemisphere (**Fig. 1A**). The same dendritic segments were imaged beginning at postnatal day (P) 28 and every four days thereafter with two additional imaging sessions in adulthood (P72 and P84) (**Fig. 1A**). Hence, the longitudinal imaging encompassed pre-, peri-, and post-pubertal stages(Laviola et al., 2003). Three animals dropped out during the study due to growth-induced prism shifts (**Supplementary Fig. S1**). For each mouse, spines on 12-15 dendritic segments (average total dendritic length of 402.2 µm ± 33.93 µm (mean ± SEM)) were analyzed (**Supplementary Fig. S1**).

**Figure 1.**
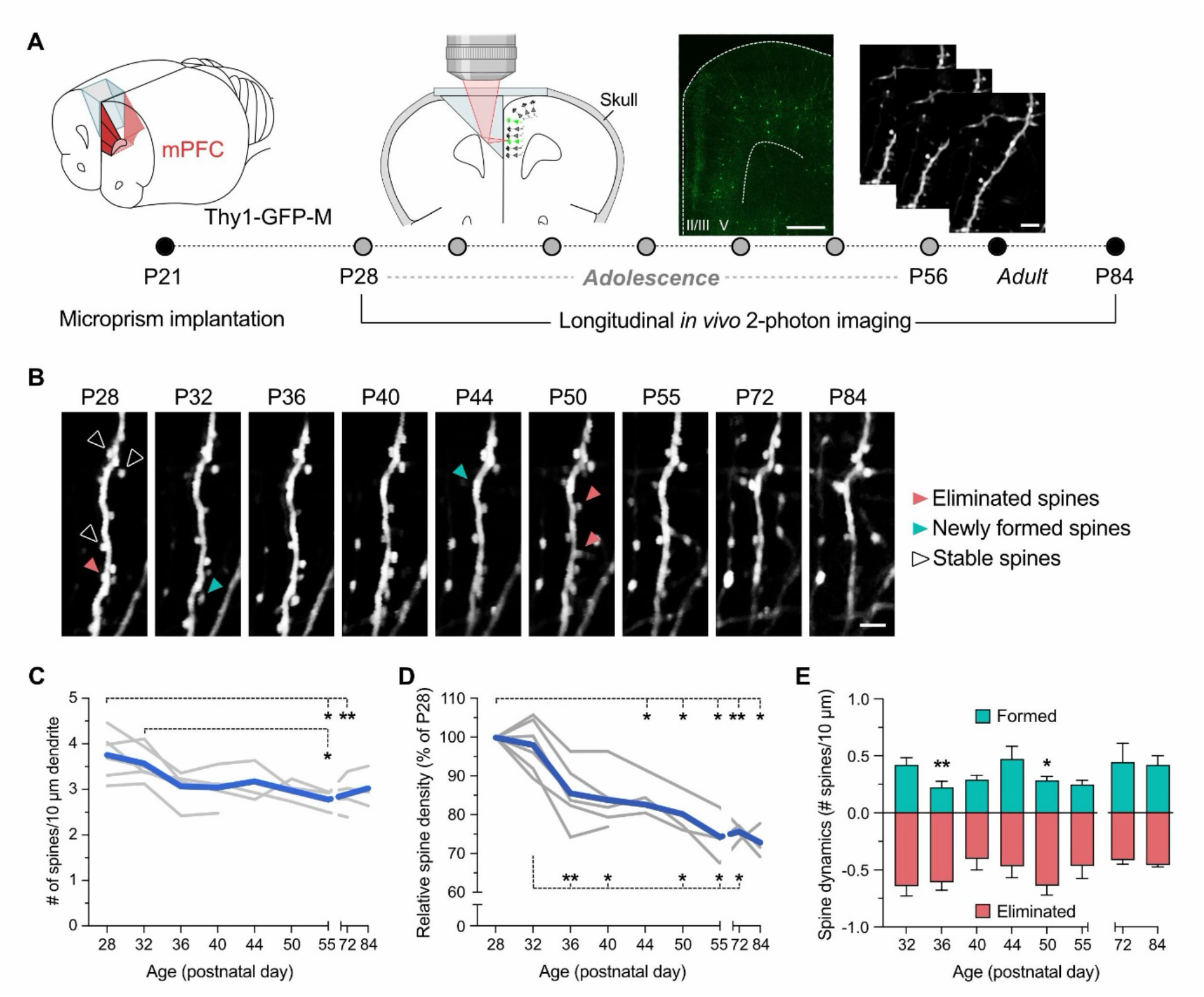
Developmental trajectory of spine dynamics across prefrontal maturation in vivo. **(A)** Schematic representation of the experimental setup. At P21 a right-angled microprism attached to a cranial window was implanted into the subdural space of Thy1-GFP-M mice. High-resolution two-photon imaging commenced at P28. Images of the same dendritic segments of layer II/III were acquired every 4 days during adolescence, once at P72 and once at P84. Scale bars = 500 μm (overview) and 5 μm (zoom-ins). **(B)** Representative two-photon images of the same dendritic segment at different imaging time points, with stable spines indicated by empty arrowheads, eliminated spines by red arrowheads, and newly formed spines by turquois arrowheads. Scale bar = 5 μm. **(C)** Number of spines per 10 μm dendrite along development. Blue line represents the overall mean, whereas the grey lines depict the mean of each animal. *p < 0.05, **p < 0.01, based on post-hoc test following mixed-effects model with a significant main effect of age. **(D)** Percent change in spine density relative to P28. Blue line represents the overall mean, whereas the grey lines depict the mean of each animal. *p < 0.05, **p < 0.01, based on post-hoc test following mixed-effects model with a significant main effect of age. **(E)** Spine dynamics assessed by comparing the number of eliminated vs newly formed spines at each maturational time point. *p < 0.05, **p < 0.01, based on post-hoc test following mixed-effects model with a significant two-way interaction between age and process. The data are means ± SEM. N = 6 mice, whereby three mice did not reach the final age (postnatal day 84) due to a shift in the microprism.

Repeated imaging of the same spines enabled quantification of synapse formation, elimination, and stabilization (**Fig. 1B**). Spine density declined significantly across prefrontal maturation (**Fig. 1C**), as indicated by a mixed-effects model demonstrating a significant main effect of age (*F*_(2.3,7.8)_ = 11.58, *p* < 0.01). Post hoc comparisons confirmed a significant reduction in absolute spine number between P28 and P55 (*p* < 0.05) or P72 (*p* < 0.01), and between P32 and P55 (*p* < 0.05) (**Fig. 1C**). Normalizing spine density to P28 yielded the same pattern (**Fig. 1D**), with a significant main effect of age (*F*_(1.8,6.2)_ = 31.07, *p* < 0.001) and post hoc comparisons revealing significant differences between P28 and P44 onward (all *p* < 0.05 or *p* < 0.01) and between P32 and subsequent ages (all *p* < 0.05 or *p* < 0.01) (**Fig. 1D**).

Across all ages, spines were both formed and eliminated (**Fig. 1E**), but elimination exceeded formation at P36 and P50 (**Fig. 1E**), reflected by a significant age × process interaction (*F*_(7,44)_ = 3.63, *p* < 0.01) and by post hoc tests confirming greater elimination at P36 (*p* < 0.01) and P50 (*p* < 0.05). Taken together, these findings show age-dependent prefrontal dendritic spine remodeling, characterized by an overall decline in spine density and discrete periods of elevated elimination.

### Developmental trajectory of excitatory synapse density across prefrontal maturation

In the mammalian brain, 80-90% of excitatory synapses are located on dendritic spines, whereas over 90% of inhibitory synapses occupy dendritic shafts, somata, or axon initial segments(Berry and Nedivi, 2017; Kwon et al., 2019). Considering the age-dependent spine remodeling observed *in vivo*, we hypothesized that excitatory synapse density would follow a similar developmental trajectory. To test this, we quantified excitatory synapse density in the mPFC using dual immunolabeling and colocalization analysis of the presynaptic vesicular glutamate transporter 1 (vGluT1) and the postsynaptic density protein Homer1(Schalbetter et al., 2022; von Arx et al., 2023). Brains were collected weekly from P21 through P56 and again at P91 (**Supplementary Fig. S2**), aligning these cross-sectional measurements with the maturational stages examined *in vivo* (**Fig. 1**). Both sexes (*N* = 6 animals per sex and age) were included. All analyses were performed in layer II/III of the anterior cingulate (AC), prelimbic (PrL), and infralimbic (IL) cortices to distinguish excitatory synapse maturation across mPFC subregions (**Fig. 2A; Supplementary Fig. S2**).

**Figure 2.**
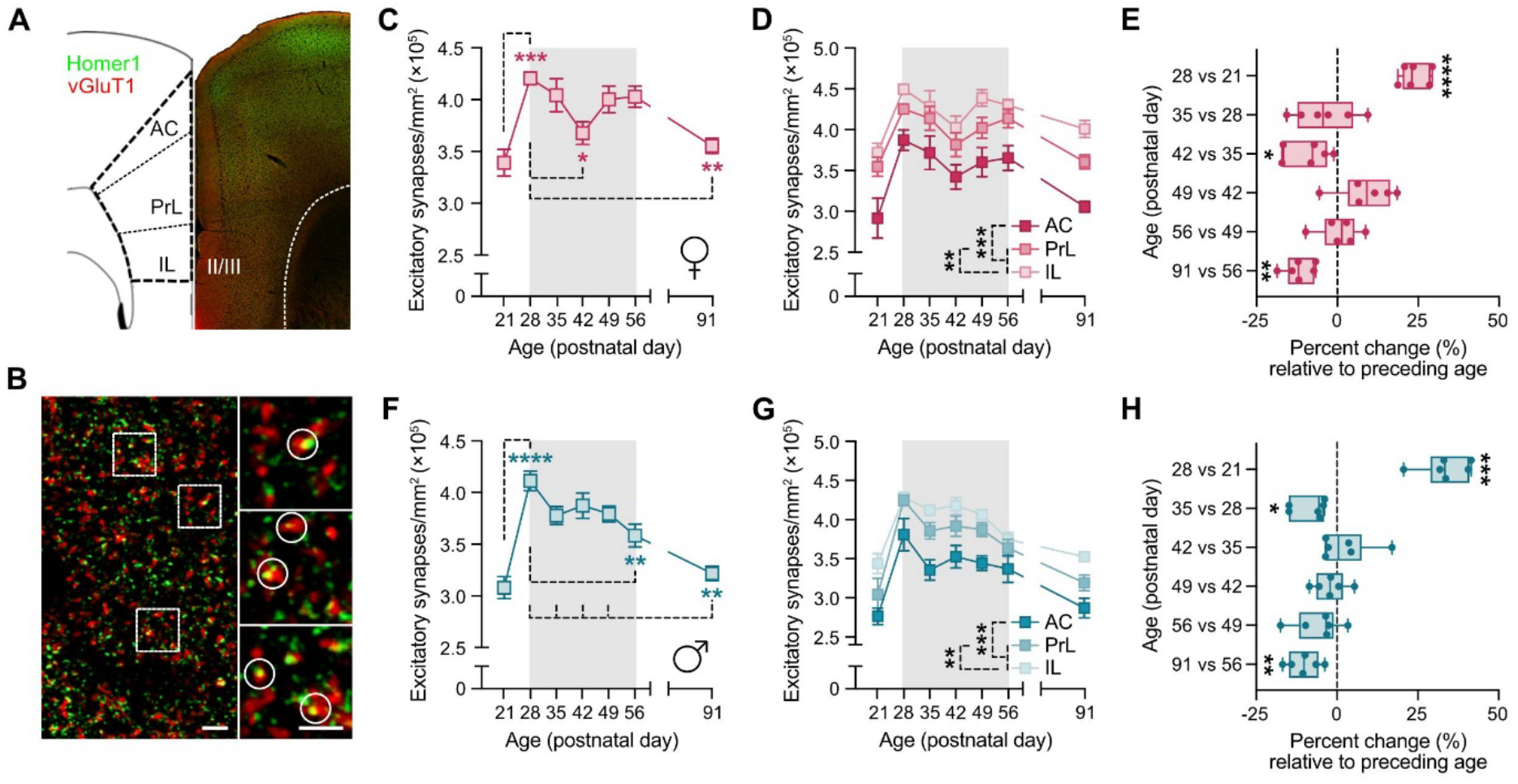
Developmental trajectory of excitatory synaptic density across prefrontal maturation. **(A)** Representative tile scan image of a brain section containing mPFC subregions (anterior cingulate (AC), prelimbic (PrL), infralimbic (IL)) stained against Homer1 (green) and vGluT1 (red). **(B)** Representative high-resolution image of layer II/III. vGluT1^+^/Homer1^+^colocalizing synapses are highlighted by white circles in magnified sections. Scale bars = 3 and 2 μm. **(C)** Densities (numbers per square millimeter) of vGluT1^+^/Homer1^+^ synapses in the mPFC across postnatal maturation in female C57BL6/N mice. \**p* < 0.05, \*\**p* < 0.01, \*\*\**p* < 0.001 based on post-hoc test following one-way ANOVA. **(D)** Densities of vGluT1^+^/Homer1^+^ synapses in prefrontal subregions across postnatal maturation in female C57BL6/N mice. \*\**p* < 0.01, \*\*\**p* < 0.001 based on post-hoc test following two-way ANOVA. **(E)** Percent change in excitatory synapse density at each postnatal age relative to the preceding age in female C57BL6/N mice. \**p* < 0.05, \*\**p* < 0.01, and \*\*\*\**p* < 0.0001, based on one-sample *t*-tests against the baseline of 0 (representing no change from the prior week). **(F)** Densities of vGluT1^+^/Homer1^+^ synapses in the mPFC across postnatal maturation in male C57BL6/N mice. \*\**p* < 0.01, \*\*\*\**p* < 0.0001 based on post-hoc test following one-way ANOVA. **(G)** Densities of vGluT1^+^/Homer1^+^ synapses in prefrontal subregions across postnatal maturation in male C57BL6/N mice. \*\**p* < 0.01, \*\*\**p* < 0.001 based on post-hoc test following two-way ANOVA. **(H)** Percent change in excitatory synapse density at each postnatal age relative to the preceding age in male C57BL6/N mice. \**p* < 0.05, \*\**p* < 0.01, and \*\*\**p* < 0.001, based on one-sample *t*-tests against the baseline of 0 (representing no change from the prior week). For C,D,F,G, the data are means ± SEM. *N* = 6 mice per group and age.

vGluT1 and Homer1 puncta were abundant across subregions (**Fig. 2A**). Automated colocalization analysis of vGluT1^+^/Homer1^+^ puncta (**Fig. 2B**) revealed a significant main effect of age in both sexes (females: *F*_(6, 35)_ = 7.107, *p* < 0.0001; males: *F*_(6, 35)_ = 14.35, *p* < 0.0001). In either sex, the number of vGluT1^+^/Homer1^+^ synapses markedly increased between P21 and P28 (*p* < 0.001 and *p* < 0.0001), followed by a decline across adolescence (**Fig. 2C,F**). In males, this decline was gradual, yielding significant reductions between P28 and P56 (*p* < 0.01) and P91 (*p* < 0.01) (**Fig. 2F**). In females, the trajectory was biphasic: after the initial increase to P28, excitatory synapse density decreased through P42 (*p* < 0.05), rebounded between P49-P56, and then dropped again significantly by P91 (*p* < 0.01; **Fig. 2C**). In either sex, this pattern was consistent across all mPFC subregions (**Fig. 2D,G**), although the number of vGluT1^+^/Homer1^+^ synapses differed across AC, PrL, and IL subregions. Across ages and in both sexes, excitatory synapse density was highest in IL, intermediate in PrL, and lowest in AC, as supported by significant main effects of subregion (females: *F*_(2,105)_ = 48.38, *p* < 0.0001; males: *F*_(2,105)_ = 44.84, *p* < 0.0001) and age (females: *F*_(6,105)_ = 13.88, *p* < 0.0001; males: *F*_(6,105)_ = 28.79, *p* < 0.0001). (**Fig. 2D,G**). Collectively, these findings show that excitatory synapses in the mPFC undergo age-dependent, regionally conserved remodeling across postnatal maturation, characterized by an early juvenile increase followed by a gradual decline in males and a biphasic decline in females across adolescence into adulthood.

To further monitor the weekly progression, we calculated the percent change in excitatory synapse density at each postnatal age relative to the preceding age. The percent-change values were then compared to the baseline of 0 (representing no change from the prior week) using one-sample *t*-tests. The largest increases occurred between P21 and P28 in both sexes, with females showing a 20-30 % rise (*t*_(5)_ = 13.9, *p* < 0.0001; **Fig. 2E**) and males a 25-40 % rise (*t*_(5)_ = 10.9, *p* < 0.001; **Fig. 2H**). Following this peak, females displayed significant declines between P35 and P42 (*t*_(5)_ = 3.4, *p* < 0.05) and between P56 and P91 (*t*_(5)_ = 6.5, *p* < 0.01) (**Fig. 2E**). In males, significant reductions were observed between P28 and P35 (*t*_(5)_ = 3.8, *p* < 0.05) and between P56 and P91 (*t*_(5)_ = 5.3, *p* < 0.01) (**Fig. 2H**). Together, these findings further show that excitatory synapse density is dynamically regulated across postnatal development, with peak increases in early adolescence followed by gradual, sex-specific reductions.

### Developmental trajectory of inhibitory synapse density across prefrontal maturation

We next investigated inhibitory synapse development in the mPFC using colocalization analysis of the presynaptic vesicular GABA transporter (vGAT) and the postsynaptic marker Gephyrin(Schalbetter et al., 2022) (**Fig. 3B**).

**Figure 3.**
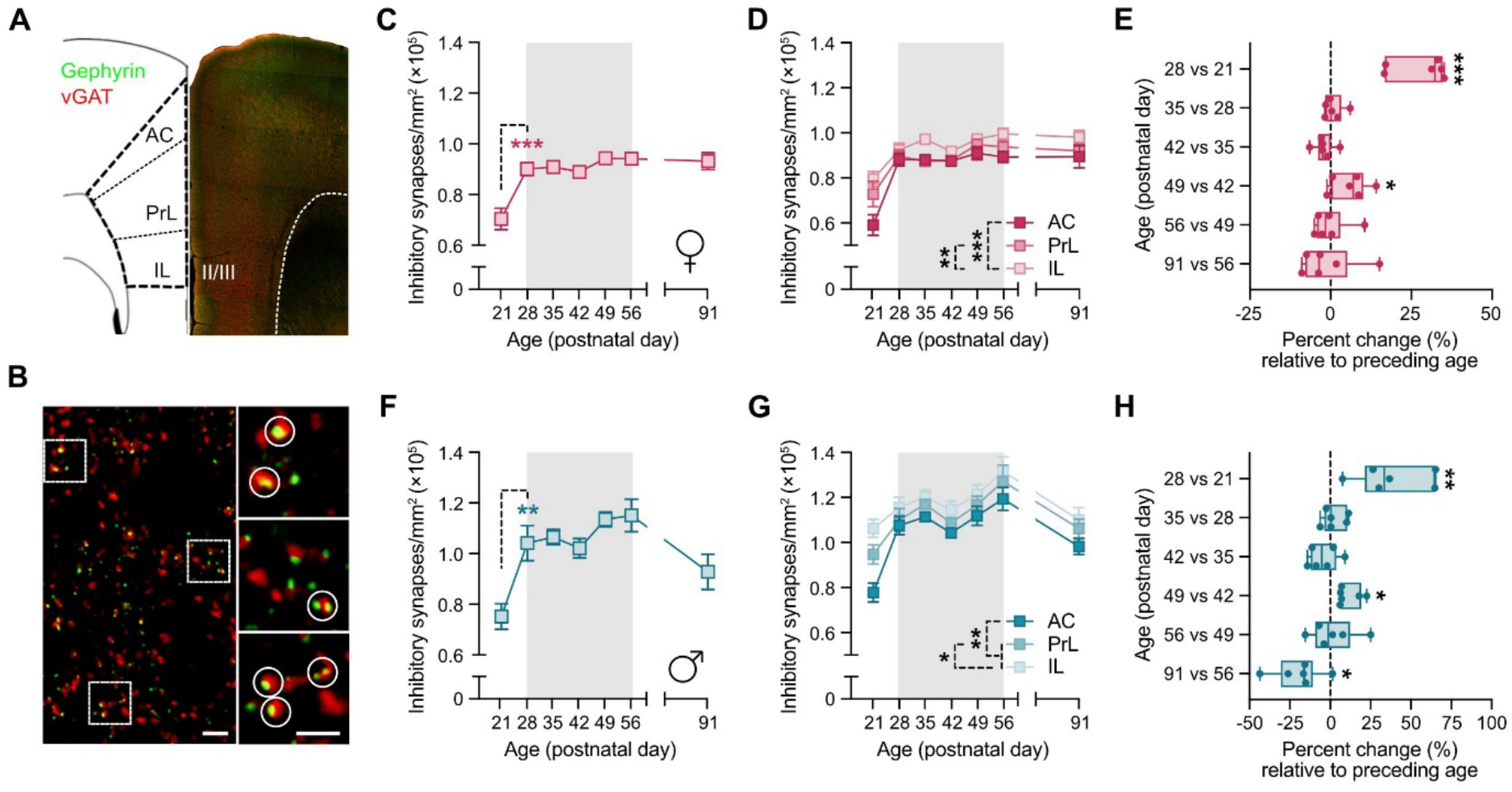
Developmental trajectory of inhibitory synaptic density across prefrontal maturation. **(A)** Representative tile scan image of a brain section containing mPFC subregions (anterior cingulate (AC), prelimbic (PrL), infralimbic (IL)) stained against Gephyrin (green) and vGAT (red). **(B)** Representative high-resolution image of layer II/III. vGAT^+^/Gephyrin^+^ colocalizing synapses are highlighted by white circles in magnified sections. Scale bars = 3 and 2 μm. **(C)** Densities (numbers per square millimeter) of vGAT^+^/Gephyrin^+^ synapses in the mPFC across postnatal maturation in female C57BL6/N mice. \*\*\**p* < 0.001 based on post-hoc test following one-way ANOVA. **(D)** Densities of vGAT^+^/Gephyrin^+^ synapses in prefrontal subregions across postnatal maturation in female C57BL6/N mice. \*\**p* < 0.01, \*\*\**p* < 0.001 based on post-hoc test following two-way ANOVA. **(E)** Percent change in inhibitory synapse density at each postnatal age relative to the preceding age in female C57BL6/N mice. \**p* < 0.05, \*\**p* < 0.01, and \*\*\*\**p* < 0.0001, based on one-sample *t*-tests against the baseline of 0 (representing no change from the prior week). **(F)** Densities of vGAT^+^/Gephyrin^+^ synapses in the mPFC across postnatal maturation in male C57BL6/N mice. \*\**p* < 0.01 based on post-hoc test following one-way ANOVA. **(G)** Densities of vGAT^+^/Gephyrin^+^ synapses in prefrontal subregions across postnatal maturation in male C57BL6/N mice. \**p* < 0.05, \*\**p* < 0.01 based on post-hoc test following two-way ANOVA. **(H)** Percent change in inhibitory synapse density at each postnatal age relative to the preceding age in male C57BL6/N mice. \**p* < 0.05, \*\**p* < 0.01, and \*\*\*\**p* < 0.0001, based on one-sample *t*-tests against the baseline of 0 (representing no change from the prior week). For C,D,F,G, the data are means ± SEM. *N* = 6 mice per group and age.

Abundant vGAT and Gephyrin puncta were observed across the regions of interest (**Fig. 3A**). As with excitatory synapses, inhibitory vGAT^+^/Gephyrin^+^ synapses increased significantly from P21 to P28 in both sexes (**Fig. 3C,F**). However, unlike the developmental trajectory of excitatory synapses (**Fig. 2**), no significant changes were observed throughout adolescence. The number of vGAT^+^/Gephyrin^+^ synapses remained largely stable between P28 and P91 in females, whereas some minor non-significant fluctuations occurred in males (**Fig. 3C,F**). These findings were supported by a significant main effect of age (females: *F*_(6,35)_ = 10.49, *p* < 0.0001; males: *F*_(6,35)_ = 6.63, *p* < 0.0001), with post hoc analyses confirming the significant increase in inhibitory synapses between P21 and P28. In both sexes, this developmental pattern was consistent across all mPFC subregions (**Fig. 3D,G**). Across ages and in both sexes, inhibitory synapse density was highest in IL, intermediate in PrL, and lowest in AC, as supported by significant main effects of subregion (females: *F*_(2,105)_ = 16.45, *p* < 0.0001; males: *F*_(2,105)_ = 15.39, *p* < 0.0001) and age (females: *F*_(6,105)_ = 24.21, *p* < 0.0001; males: *F*_(6,105)_ = 17.63, *p* < 0.0001). Together, these data show that inhibitory synapse density in the mPFC is largely stable across adolescence, with a conserved subregional pattern in both sexes.

Consistent with excitatory synapses (**Fig. 2**), largest week-to-week changes in inhibitory synapse density occurred between P21 and P28 in both sexes, with females exhibiting a 15-35 % rise (*t*_(5)_ = 7.8, *p* < 0.001; **Fig. 3E**) and males a 10-65 % rise (*t*_(5)_ = 7.8, *p* < 0.001; **Fig. 3H**). Beyond this early adolescent peak, inhibitory synapse density remained largely stable in females, with a small but significant (*t*_(5)_ = 2.6, *p* < 0.05) increase occurring between P42 and P49 (**Fig. 3E**). Males showed a similar P42-P49 increase (*t*_(5)_ = 3.8, *p* < 0.05), followed by a decline between P56 and P91 (*t*_(5)_ = 3.2, *p* < 0.05; **Fig. 3H**). Overall, inhibitory synapse density in the mPFC is established rapidly by early adolescence and remains relatively stable thereafter with only modest, sex-specific adjustments during later maturation.

### Developmental trajectory of the excitatory/inhibitory balance across prefrontal maturation

Based on the analyses of excitatory (**Fig. 2**) and inhibitory (**Fig. 3**) synapse densities, we next calculated an excitatory/inhibitory (E/I) ratio at each age by dividing the number of excitatory puncta by the number of inhibitory puncta. The E/I ratio undergoes marked age-dependent changes across postnatal development in both sexes (**Fig. 4**), as supported by a significant main effect of age (females: *F*_(6,35)_ = 4.45, *p* < 0.01; males: *F*_(6,35)_ = 2.83, *p* < 0.05). In females, the E/I ratio was highest at P21 and progressively declined across adolescence, reaching its minimum at P91 (P21 vs. P91: *p* < 0.001; P28 vs. P91: *p* < 0.05; **Fig. 4A**). In males, the E/I ratio similarly peaked at P21 and decreased gradually, reaching its lowest point at P56 (P21 vs. P56: *p* < 0.05; **Fig. 4B**). These findings indicate that prefrontal circuit maturation involves a developmental shift toward lower E/I ratios from adolescence into adulthood, with partially distinct temporal trajectories in females and males.

**Figure 4.**
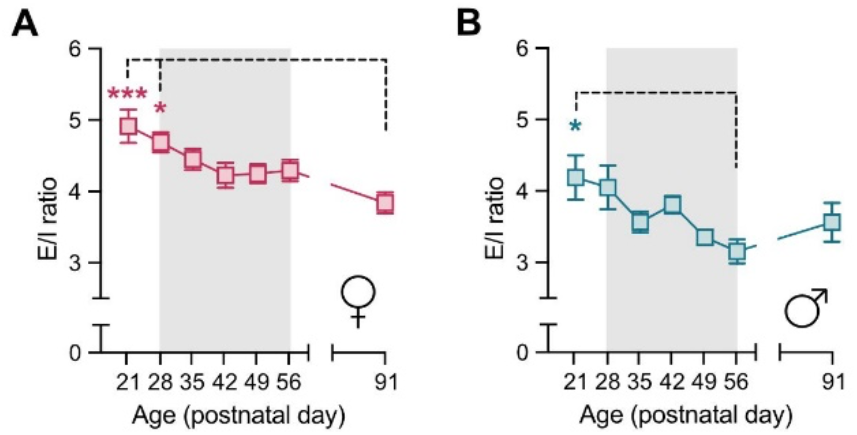
Developmental trajectory of excitatory/inhibitory (E/I) balance across prefrontal maturation. **(A)** E/I ratio across postnatal maturation in female C57BL6/N mice. \*\*\**p* < 0.001, \**p* < 0.05 based on post-hoc test following one-way ANOVA. **(B)** E/I ratio across postnatal maturation in male C57BL6/N mice. \**p* < 0.05 based on post-hoc test following one-way ANOVA. All data are means ± SEM. *N* = 6 mice per group and age.

### Developmental trajectory of pan-presynaptic density across prefrontal maturation

Bassoon-positive presynaptic terminals provided a pan-presynaptic measure of mPFC maturation that complements the excitatory (vGluT1/Homer1) and inhibitory (vGAT/Gephyrin) analyses. Bassoon, an active-zone scaffolding protein present at both excitatory and inhibitory synapses, is involved in vesicle docking, release, and synaptic protein organization and marks functional presynaptic terminals(Richter et al., 1999; Gundelfinger et al., 2015). Quantification of Bassoon^+^ puncta across juvenile to adult stages (**Fig. 5A, B, Supplementary Fig. S2)** revealed clear age-dependent changes in both sexes, supported by a significant main effect of age (females: *F*_(6,55)_ = 30.34, *p* < 0.0001; males: *F*_(6,54)_ = 25.41, *p* < 0.0001)

Bassoon density increased sharply between P21 and P28 in females and males (*p*’s < 0.0001; **Fig. 5C,F**), mirroring the early rise in excitatory and inhibitory synapses. This early increase was followed by a gradual decline across adolescence, with differences between sexes. Females exhibited a significant reduction between P28 and P49 (*p* < 0.05; **Fig. 5C**), whereas males showed a significant reduction between P28 and P56, as well as P35 and P56 (*p* < 0.05; **Fig. 5F**). In either sex, this developmental pattern was consistent across AC, PrL, and IL (**Fig. 5D,G**), although absolute Bassoon density differed by subregion. being highest in IL, intermediate in PrL, and lowest in AC (**Fig. 5D,G**) as supported by significant main effects of subregion (females: *F*_(2,165)_ = 66.77, *p* < 0.0001; males: *F*_(2,162)_ = 19.52, *p* < 0.0001) and age (females: *F*_(6,165)_ = 62.36, *p* < 0.0001; males: *F*_(6,162)_ = 54.91, *p* < 0.0001).

**Figure 5.**
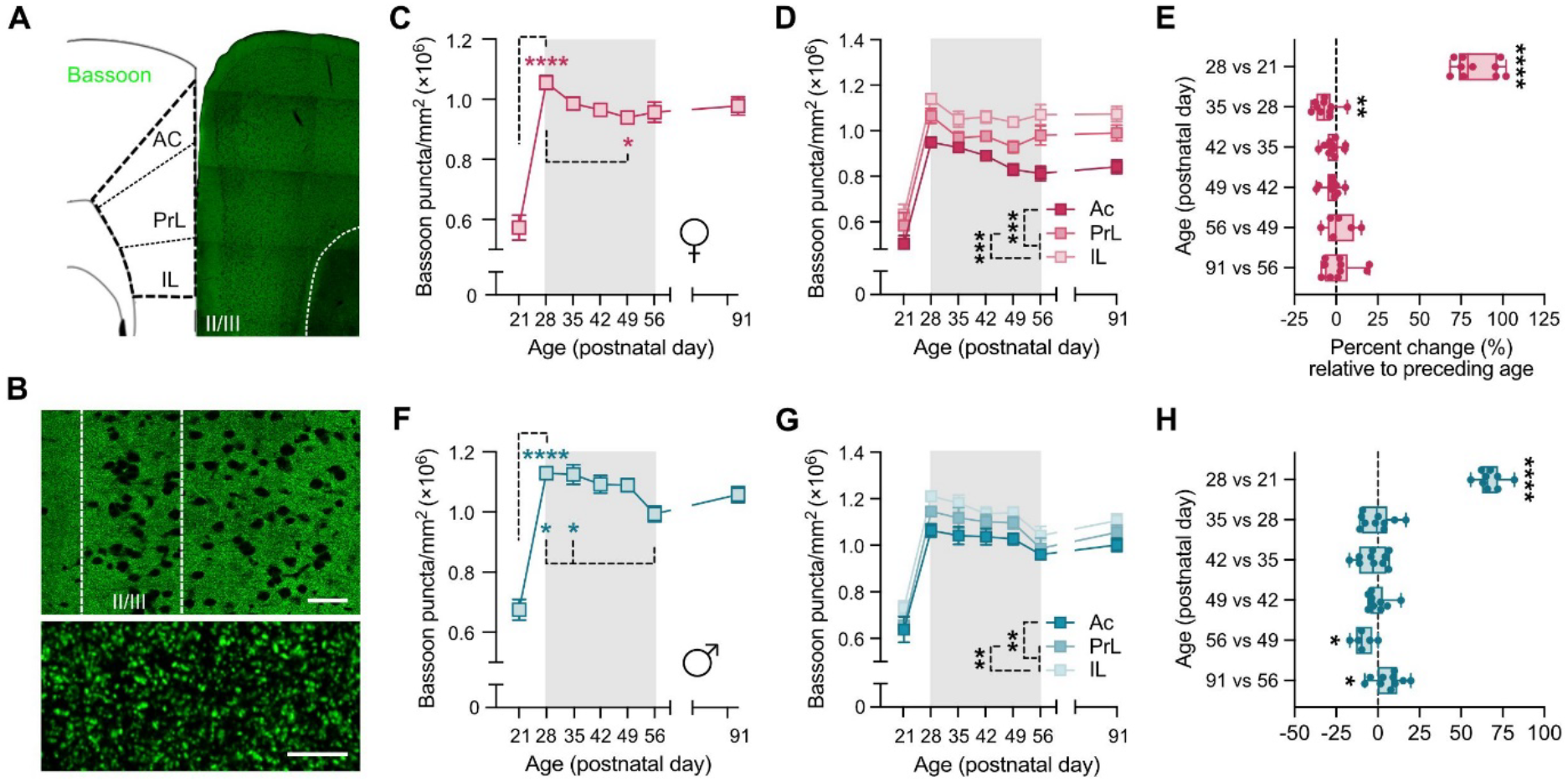
Developmental trajectory of pan-presynaptic density across prefrontal maturation. **(A)** Representative tile scan image of a brain section containing mPFC subregions (anterior cingulate (AC), prelimbic (PrL), infralimbic (IL)) stained against Bassoon. **(B)** Representative high-resolution images of layer II/III (highlighted by the dashed line) at different magnifications. Scale bars = 50 and 5 μm. **(C)** Densities (numbers per square millimeter) of Bassoon^+^ presynaptic puncta in the mPFC across postnatal maturation in female C57BL6/N mice. \**p* < 0.05, \*\*\*\**p* < 0.0001 based on post-hoc test following one-way ANOVA. **(D)** Densities of Bassoon^+^ presynaptic puncta in prefrontal subregions across postnatal maturation in female C57BL6/N mice. \*\*\**p* < 0.001 based on post-hoc test following two-way ANOVA. **(E)** Percent change in pan-presynaptic density at each postnatal age relative to the preceding age in female C57BL6/N mice. \*\**p* < 0.01 and \*\*\*\**p* < 0.0001, based on one-sample *t*-tests against the baseline of 0 (representing no change from the prior week). **(F)** Densities of Bassoon^+^ presynaptic puncta in the mPFC across postnatal maturation in male C57BL6/N mice. \**p* < 0.05, \*\*\*\**p* < 0.0001 based on post-hoc test following one-way ANOVA. **(G)** Densities of Bassoon^+^ presynaptic puncta in prefrontal subregions across postnatal maturation in male C57BL6/N mice. \*\**p* < 0.01 based on post-hoc test following two-way ANOVA. **(H)** Percent change in pan-presynaptic density at each postnatal age relative to the preceding age in male C57BL6/N mice. \**p* < 0.05 and \*\*\*\**p* < 0.0001, based on one-sample *t*-tests against the baseline of 0 (representing no change from the prior week). For C,D,F,G, the data are means ± SEM. *N* = 10 mice per group and age except for male P28, where *N* = 9 and for P21 and P56 where *N* = 6 (for both sexes).

Consistent with excitatory and inhibitory synapses, the week-to-week analysis revealed that the largest increase in Bassoon density occurred between P21 and P28 in both sexes, with females exhibiting a 70-100 % rise (*t*_(9)_ = 21.0, *p* < 0.0001; **Fig. 5E**) and males a 55-80 % rise (*t*_(8)_ = 27.2, *p* < 0.0001; **Fig. 5H**). After this peak, females showed a significant decline between P28 and P35 (*t*_(9)_ = 3.4, *p* < 0.01; **Fig. 5E**). Males exhibited a significant reduction between P49 and P56 (*t*_(5)_ = 3.7, *p* < 0.05; **Fig. 5H**), followed by a small but significant increase between P56 and P91 (*t*_(9)_ = 2.4, *p* < 0.05; **Fig. 5H**). Together, these analyses indicate that presynaptic density in the mPFC is established rapidly during early adolescence and subsequently refined in a sex-specific manner, with earlier declines in females than males.

### Developmental trajectory of microglia-mediated synaptic pruning across prefrontal maturation

Synaptic remodeling across postnatal brain maturation is shaped in part by microglia, the resident immune cells of the brain parenchyma(Paolicelli et al., 2011; Schafer et al., 2012; Scott-Hewitt et al., 2020; Faust et al., 2025). In addition to their classical immunological functions(Wolf et al., 2017), microglia can promote synaptic pruning, a process in which superfluous synapses are eliminated via direct phagocytosis(Paolicelli et al., 2011; Schafer et al., 2012; Scott-Hewitt et al., 2020; Faust et al., 2025). Given the adolescent periods of elevated spine elimination observed *in vivo* (**Fig. 1**), we hypothesized that microglial engulfment of synaptic elements accompanies the age-dependent fluctuations. To test this hypothesis, we profiled the developmental trajectory of microglial engulfment of presynaptic elements in the mPFC across the same juvenile to adult stages (**Supplementary Fig. S2**) by calculating a phagocytic index based on Bassoon^+^ puncta contained within surface-rendered Iba1^+^ microglia (**Fig. 6A**).

**Figure 6.**
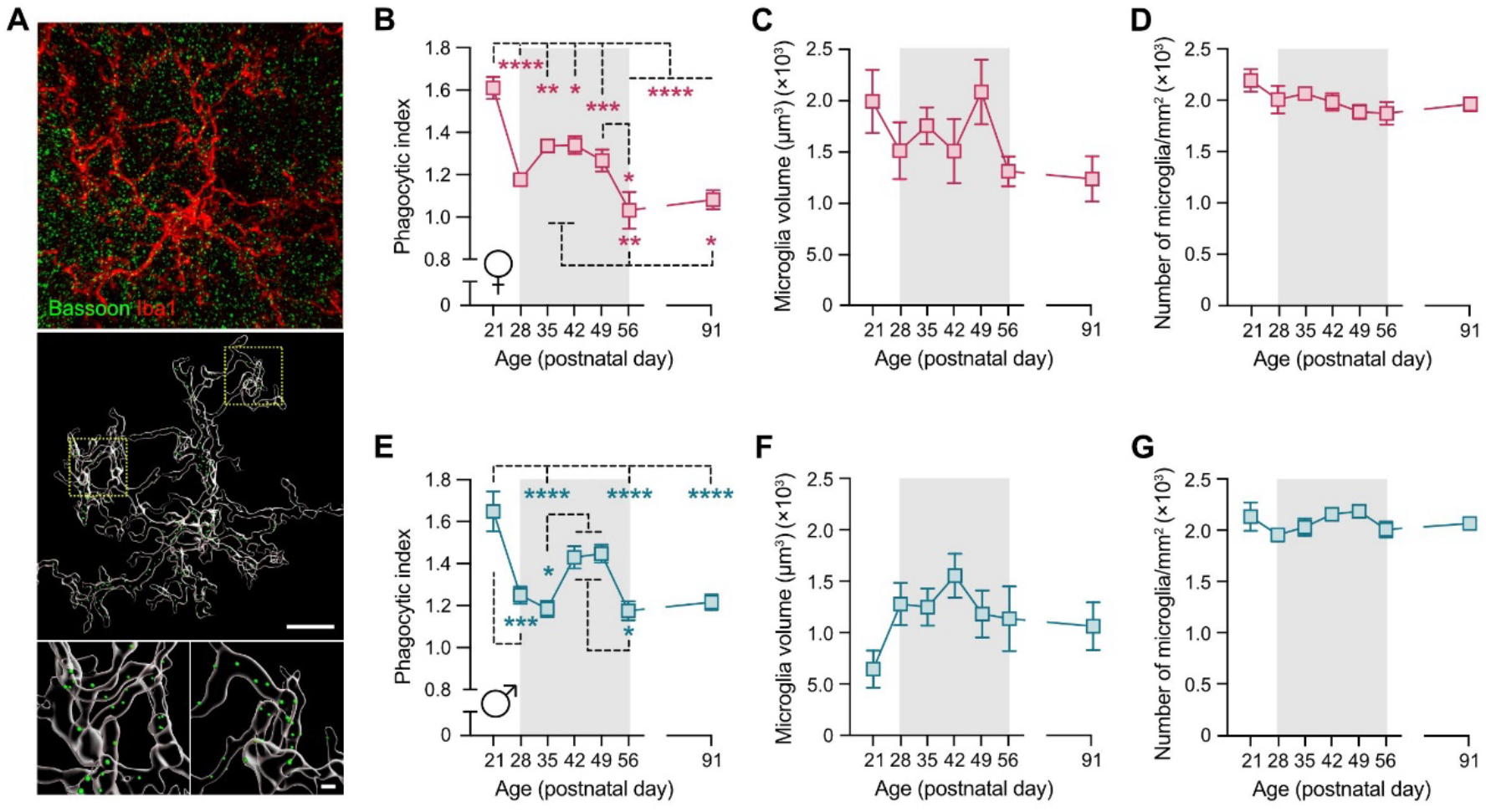
Developmental trajectory of microglia-mediated synaptic pruning across prefrontal maturation. **(A)** The photomicrograph shows a representative z-stack image of a double-immunofluorescence stain against Iba1 (red) as microglial marker and Bassoon (green) as pan-presynaptic marker before (top) and after (bottom) surface rendering and reconstruction with Imaris image analysis software. Bassoon^+^ presynaptic puncta within microglia appear as green dots in the reconstructed image. Scale bars = 10 and 1 μm. **(B)** Phagocytic index (Bassoon^+^ presynaptic puncta within microglia) in the mPFC across postnatal maturation in female C57BL6/N mice. \**p* < 0.05, \*\**p* < 0.01, \*\*\**p* < 0.001, \*\*\*\**p* < 0.0001 based on post-hoc test following one-way ANOVA. **(C)** Average microglia volume (cubic micrometer) in the mPFC across postnatal maturation in female C57BL6/N mice. **(D)** Densities (numbers per square millimeter) of microglia in the mPFC across postnatal maturation in female C57BL6/N mice. **(E)** Phagocytic index (Bassoon^+^ presynaptic puncta within microglia) in the mPFC across postnatal maturation in male C57BL6/N mice. \**p* < 0.05, \*\*\**p* < 0.001, \*\*\*\**p* < 0.0001 based on post-hoc test following one-way ANOVA. **(F)** Average microglia volume (cubic micrometer) in the mPFC across postnatal maturation in male C57BL6/N mice. **(G)** Densities (numbers per square millimeter) of microglia in the mPFC across postnatal maturation in male C57BL6/N mice. All data are means ± SEM. *N* = 6 mice per group and age except for male P28, where *N* = 5.

The phagocytic index varied across age in both females and males (**Fig. 6B,E**), as indicated by a significant main effect of age (females: *F*_(6,35)_ = 14.62, *p* < 0.0001; males: *F*_(6,34)_ = 10.70, *p* < 0.0001), following a biphasic pattern marked by a juvenile peak at P21 and a second peak during mid-adolescence (**Fig. 6B,E**). While this overall biphasic pattern was similar between sexes, the onset of the second peak occurred earlier in females than in males (**Fig. 6B,E**). In females, the decline in the phagocytic index between P21 and P28 was followed by an increase between P28 and P35, peaking at P42, and a subsequent decline thereafter (**Fig. 6B**). This trajectory led to significant differences between P21 and all other time points (*p* < 0.05 to *p* < 0.0001), as well as between P35 or P42 and P56 or P91 (*p* < 0.05 to *p* < 0.01) (**Fig. 6B**). In males, the decline in the phagocytic index continued until P35, after which it increased between P42 and P49, followed by a subsequent decline into adulthood (**Fig. 6E**) resulting in significant differences between P21 and P28, P35, P56, and P91 (*p* < 0.001 to *p* < 0.0001), between P35 and P42 or P49 (all *p* < 0.05), and between P42 or P49 and P56 (all *p* < 0.05) (**Fig. 6E**). Together, these data show that microglia in the mPFC exhibit a biphasic, age-dependent pattern of presynaptic engulfment across postnatal maturation, with early juvenile and mid-adolescent peaks that occur earlier in females than in males.

Surface-rendered microglial morphology showed no significant differences in the average volume of microglia across juvenile to adult maturation (**Fig. 6C,F**) and the density of prefrontal microglia (number/mm^2^) (**Fig. 6D,G**). Thus, developmental changes in microglia-mediated synaptic pruning occur without major changes in microglial morphology or density.

## DISCUSSION

Our study provides a systematic characterization of developmental synaptic remodeling in the mouse mPFC by combining high-resolution, longitudinal *in vivo* imaging with quantification of synaptic densities across a broad maturational time span. Spanning early adolescent (P28) to adult (P84) stages, our longitudinal *in vivo* data of spine dynamics extend previous two-photon imaging studies, spanning P24-P28 or P64-P68(Johnson et al., 2016), P30-P31 or P44-P45(Pattwell et al., 2016), and P27-P28 or P60-P61(Boivin et al., 2018). By contrasting the trajectories of excitatory and inhibitory synapses with *in vivo* spine dynamics and pan-presynaptic measures, our study further advances prior post-mortem analyses, which typically quantified either presynaptic terminals or postsynaptic spines only, while rarely distinguishing synapse types(Gourley et al., 2012; Koss et al., 2014; Drzewiecki et al., 2016; Delevich et al., 2020; Pöpplau et al., 2024). Hence, our dataset provides both specificity and generalizability across a broad developmental period, establishing a reference atlas of normal prefrontal synaptic maturation in male and female mice and offering a framework for interpreting synaptic alterations in models of developmental disruption(Hinton et al., 2019; Miller et al., 2019; Shaw et al., 2020; Benoit et al., 2022; Schalbetter et al., 2022; Pöpplau et al., 2024).

Overall, our findings align with previous studies in humans(Huttenlocher, 1979; Petanjek et al., 2011), non-human primates(Bourgeois et al., 1994; Anderson et al., 1995; Elston et al., 2009), and rodents(Gourley et al., 2012; Koss et al., 2014; Drzewiecki et al., 2016; Delevich et al., 2020; Pöpplau et al., 2024), which demonstrate a decline in prefrontal synaptic density from adolescence to adulthood. In our study, this trajectory is supported by three independent observations, namely the longitudinal *in vivo* imaging of spine dynamics, quantification of vGluT1^+^/Homer1^+^ excitatory synapses, and measurement of the pan-presynaptic marker Bassoon. In contrast, the developmental trajectory of inhibitory synapse density followed an opposite pattern. Rather than declining, inhibitory synapse density remained stable or increased slightly from juvenile to adult stages, resulting in a progressive reduction in E/I ratios across adolescence. These findings are consistent with previous electrophysiological studies indicating that inhibitory drive within prefrontal circuits strengthens during adolescence, leading to lower E/I ratios in adulthood compared with earlier developmental stages(Caballero and Tseng, 2016; Caballero et al., 2021).

The inclusion of male and female animals in our study revealed both conserved and sex-specific patterns of prefrontal circuit maturation from juvenile to adult stages. In both sexes, excitatory synapses exhibited an early juvenile increase followed by an adolescent decline, whereas inhibitory synapse density rose early during the juvenile period and remained largely stable thereafter. Despite these shared features, we observed some sex differences in the timing of synaptic maturation. Consistent with previous post-mortem analyses of pan-presynaptic markers in the PFC of rats(Drzewiecki et al., 2016), the adolescent decline in Bassoon-immunoreactive presynaptic density occurred earlier in females than in males. Moreover, excitatory synapse remodeling was biphasic in females, with a rebound in density between P49 and P56, whereas males showed a more gradual, monotonic decline. Similarly, the developmental decline in the E/I ratio occurred earlier in males, reaching its minimum at P56, while females displayed a more protracted reduction extending to P91. These findings highlight that, although the overall patterns of prefrontal synaptic maturation are conserved across sexes, the precise timing and dynamics differ, underscoring the importance of including both males and females in developmental studies.

The temporal pattern of synaptic remodeling was also mirrored in the age-dependent phagocytic activity of prefrontal microglia. Both males and females exhibited a biphasic pattern of microglial engulfment, with peaks in early juvenile and mid-adolescent stages. Interestingly, the sharp decrease in the phagocytic index between P21 and P28 corresponded to the marked rise in synaptic density during the same period, highlighting a close temporal relationship between microglia-mediated pruning and the early adolescent surge in synapse density at P28. Furthermore, the subsequent increase in the phagocytic index across mid-to-late adolescence corresponded to a phase in which the density of the pan-presynaptic marker Bassoon and vGluT1^+^/Homer1^+^ excitatory synapses declined. Together, although correlative in nature, these data provide a high-resolution dataset showing the temporal association between microglia-mediated engulfment of synaptic elements and corresponding alterations in synaptic densities. Notably, consistent with the partly sex-dependent developmental trajectories of synaptic densities, the phagocytic index was also influenced by sex. The second peak of microglial engulfment occurred earlier in females than in males, paralleling the observation that the density of Bassoon and vGluT1^+^/Homer1^+^ excitatory synapses declined earlier in adolescence in females than in males. These findings add to the emerging literature demonstrating notable sex differences in microglial biology in both health and disease(Schwarz and Bilbo, 2012; Hanamsagar et al., 2017; Guneykaya et al., 2018; VanRyzin et al., 2019), suggesting that microglial influence on PFC maturation may be sexually dimorphic. It should be noted, however, that in addition to microglia-mediated synaptic pruning, other mechanisms are likely to contribute to synaptic remodeling during prefrontal maturation(Hall and Bray, 2022), including microglia-associated non-phagocytic processes(Weinhard et al., 2018; Cheadle et al., 2020), as well as synaptic pruning by astrocytes or oligodendrocyte precursor cells(Chung et al., 2013; Auguste et al., 2022).

Although we do not have a parsimonious explanation for the pronounced increase in synaptic density between P21 and P28, it is noteworthy that this interval coincides with the post-weaning period of the animals. During this time, juvenile mice are separated from their rearing mothers and relocated to new home cages. To our knowledge, no study has explicitly addressed the influence of weaning on synaptic remodeling. However, the process of weaning and relocation to a novel cage environment is likely associated with exposure to a variety of novel sensory and social stimuli, which may promote the formation of new synapses and/or the strengthening of existing ones. This may be reflected in the observed sharp increase in synaptic density during the transition from the juvenile to the early adolescent phase, consistent with evidence that sensory experience can both attenuate synaptic pruning and promote synaptic strengthening(Poeggel et al., 2003; Cheadle et al., 2020; Faust et al., 2021). We acknowledge several limitations of our study. First, because the study design focused on combining high-resolution longitudinal *in vivo* imaging with quantification of synaptic densities across a broad maturational time span, it did not include specific experimental manipulations that would allow testing causal relationships. Nevertheless, even if our dataset is descriptive, it provides a comprehensive quantitative atlas of normative prefrontal synaptic maturation in male and female mice, which can readily aid the interpretation of synaptic alterations reported in previous or future studies of developmental perturbations in mouse models. Second, while the initial *in vivo* imaging is unique in providing high-resolution measurements of spine dynamics in the mPFC, the dropout of several male animals prevented us from fully assessing potential sex differences in these longitudinal measurements. However, this limitation does not undermine the overall validity of the *in vivo* imaging dataset, as subsequent quantification of all other synaptic measures and microglial phagocytic activity was performed using a sufficient number of male and female mice.

Notwithstanding these limitations, our study shows that prefrontal synaptic maturation in mice is a temporally coordinated, partly sexually dimorphic process. Excitatory and inhibitory synapses, together with microglial phagocytic activity, follow distinct yet interrelated developmental trajectories from the juvenile to adult stages. By creating a detailed reference atlas of normative synaptic and microglial remodeling, our work offers a valuable resource for the neuroscience community. It provides a critical foundation for future studies aimed at identifying, contextualizing, and ultimately mitigating synaptic disruptions in models of neurodevelopmental disorders.

## MATERIAL AND METHODS

### Animals

Male and female C57BL6/N mice were used for the cross-sectional analyses of synaptic density along adolescence. Female and male C57BL6/N breeder mice were obtained from Charles River Laboratories (Sulzfeld, Germany). Longitudinal *in vivo* imaging experiments were conducted in hemizygous transgenic male and female Thy1-GFP-M mice (Thy1-GFP-M line, JAX: 007788, The Jackson Laboratory, Bar Harbor, USA). All animals were housed in groups of 2-5 per individually ventilated cage (IVCs; Allentown Inc., Bussy-Saint-Georges, France) in a temperature-and humidity-controlled (21°C ± 3°C, 50 ± 10%) specific pathogen-free holding room under reversed light conditions. All animals had *ad libitum* access to standard rodent chow food (Kliba 3336, Kaiseraugst, Switzerland) and water. All procedures were previously approved by the Cantonal Veterinarian’s Office of Zurich. Every effort was made to minimize the number of animals and their suffering. The number of animals used in each experiment is specified in the legends of the figures and supplementary figures.

### Timed-breeding procedure

To produce C57BL6/N and hemizygous Thy1-GFP-M offspring of specific ages for subsequent immunohistochemical and *in vivo* imaging experiments, respectively, a timed-mating procedure was used as previously described(Notter et al., 2018; Schaer et al., 2025). To this end, female and male mice were paired using a 2:1 ratio. Mating was verified by the presence of a vaginal plug. Two females with verified mating were housed together until weaning of the offspring (postnatal day (P) 21). For the collection of post-mortem tissue, parental mice were paired on a weekly schedule, allowing all offspring to be sacrificed on the same day at their respective ages (P21, P28, P35, P42, P49, P56, and P91). To produce hemizygous Thy1-GFP-M offspring, homozygous Thy1-GFP-M male mice were paired with C57BL6/N female mice.

### Genotyping of Thy1-GFP-M offspring

Genotypes were verified at P14 by PCR using the following primers: CGG TGG TGC AGA TGA ACT T (transgene reverse, #15731, The Jackson Laboratory, Bar Harbor, USA), ACA GAC ACA CAC CCA GGA CA (transgene forward, #16072, The Jackson Laboratory, Bar Harbor, USA), and the internal positive controls CTA GGC CAC AGA ATT GAA AGA TCT (forward, #oIMR7338, The Jackson Laboratory, Bar Harbor, USA), and GTA GGT GGA AAT TCT AGC ATC ATC C (reverse, #oIMR733, The Jackson Laboratory, Bar Harbor, USA). The PCR protocol included an initial denaturation at 95°C for 3.15 minutes, followed by annealing at 60°C for 15 seconds and extension at 72°C for 20 seconds. The first cycle was followed by 35 subsequent cycles, each consisting of 15 seconds of melting at 95°C, 15 seconds at 60°C (annealing), and 20 seconds at 72°C (extension). A final extension at 72°C for 1 minute was performed. PCR products were run on a 2% agarose gel (10.5 µl per lane) in TBE buffer (Tris base 1 M, boric acid 1 M, EDTA disodium salt 0.02 M) at 120 V and visualized under UV light using RedSafe™ Nucleic Acid Staining Solution (INT-21141-1ML).

### Longitudinal in vivo two-photon imaging of dendritic spines

#### Microprism cranial window assembly

A custom-made right-angled microprism (1.5-mm side length and 1-mm width, S-BSL7, protected aluminum coating on hypotenuse surface to enable internal reflection; Optosigma) was bonded to a circular glass window (3 mm diameter coverslip) using UV-curing optical adhesive (Norland #81) on the day of the surgery.

#### Microprism implantation

The surgery for the intracranial microprism implantation was performed using methods established and validated before(Low et al., 2014). A total of 5 female and 5 male hemizygous Thy1-GFP-M mice were generated for microprism implantation. At P21, mice were anesthetized with midazolam (5 mg/kg), fentanyl (0.05 mg/kg), and medetomidine (0.5 mg/kg) anesthesia. First, a chrome steel head plate was implanted. In brief, anesthetized animals were fixed in a stereotaxic frame and the skin above the midline was carefully cut and removed. The bone was cleaned, and a bonding agent (Prime & Bond) was applied to the skull and polymerized with blue light. A round head plate was attached to the exposed bone using light-cured dental cement (Tetric EvoFlow). The skull over the PFC was left exposed for the craniotomy. Next, the craniotomy was performed using a dental drill (rotate, H-4-002). The skull was first thinned and carefully removed in small bone fragments, leaving the dura intact. At this stage, 4 males had to be excluded due to bridging veins that precluded prism implantation(Low et al., 2014). For the remaining 6 mice, an incision in the dura along the side of the sinus was created where the microprism was implanted. The microprism was then gently inserted into the subdural space within the fissure so that the prism surface sat flush opposite the hemisphere to be imaged (with the cerebral falx between the microprism surface and PFC cortex to be imaged). The area beneath the microprism (i.e., the medial portion of the PFC of the contralateral site, which was not imaged) was compressed but remained intact. Dental cement was then used to secure the glass cover slip in place. After completion of the surgery, the animals were placed in a temperature-controlled chamber until complete recovery from anesthesia after which they were placed back in their home cage and closely monitored for three consecutive days.

#### In vivo two-photon imaging in anesthetized animals

All images were acquired using a custom-made two-photon laser scanning microscope(Mayrhofer et al., 2015). The same dendritic segments were imaged in anesthetized mice (1-1.5% isoflurane) starting from P28 ± 1 and every four days thereafter until P55 ± 1. An additional imaging session was conducted at P72 (early adulthood) and P84 (adult). Anesthesia was induced with 5% isoflurane, after which the mice were transferred to the two-photon microscope where they were head-fixed. Anesthesia was maintained with 1-1.5% isoflurane, whereby depth of anesthesia was monitored and body temperature maintained using the MARTA monitoring system (Vigilitech AG, Switzerland). An eye lubricant (Vitamin A, Bausch + Lomb) was applied to prevent corneal drying during imaging. The window was cleaned with deionized water prior to imaging. Images were acquired using a water-immersed 60 × objective (NA 1, Nikon Instruments, MRD07620) with a working distance of 2.8 mm. In each imaging session, z-stacks with a resolution of 1024 × 1024 pixels (pixel size: 0.08286 x 0.08286 µm, z-step: 0.8 µm, optical zoom: 5) were acquired at a distance of 150–200 µm from the tissue surface. 5 z-stacks were acquired per animal and imaging session. The same dendritic segments were relocated at micrometer precision across imaging sessions using reference points at the edge of the microprism and coordinate systems. Due to continued skull growth between P21 and P84, some animals dropped out during the study because a shift in the microprism resulted in image loss.

#### Image analysis

Spine dynamics were analyzed using the ImageJ software. Gaussian filter and background subtraction were applied to increase signal to noise. Spines on different dendritic segments (10 – 13 dendritic segments per mouse) were then counted manually by reviewing the z-stack using previously defined criteria(Holtmaat et al., 2009; Phoumthipphavong et al., 2016). Spine dynamics were assessed by comparing the respective spines (image 2) with those from the previous imaging day (image 1). The total number of spines, the number of eliminated (present in image 1 and absent in image 2) and the number of newly formed (absent in image 1 and present in image 2) spines per 10 µm of dendrite were calculated (average total length of dendrite analyzed per animal was 402.2 µm ± 33.93 µm (mean ±SEM)). Relative spine density was calculated by dividing the number of spines/10 µm of dendrite per given time point with the number of spines/10 µm of dendrite at P28 × 100.

### Cross-sectional analyses of synaptic density using immunohistochemistry

Sample collection, immunofluorescence staining and image acquisition were performed according to established protocols(Notter et al., 2014; Notter et al., 2018; Schalbetter et al., 2022). Tissue was collected from C57BL/6N male and female mice at P21, P28, P35, P42, P49, P56, and P91, spanning preadolescent, adolescent, and adult developmental stages(Laviola et al., 2003) (**Supplementary Fig. 2**). These time points were selected to achieve high temporal resolution of synaptic remodeling across adolescence(Drzewiecki et al., 2016) and to capture potential sex-dependent differences related to pubertal onset(Drzewiecki et al., 2016; Delevich et al., 2020), which occurs between P30–P32 in females and P35–P40 in males(Nelson et al., 1990; Chowdhury et al., 2013).

Sample collection

Animals were deeply anesthetized with an overdose of pentobarbital (Esconarkon ad us. vet., Streuli Pharma AG, Switzerland) and perfused transcardially with artificial cerebrospinal fluid (aCSF, pH 7.4) with a perfusion rate of 20 ml/min. The brains were immediately removed from the skull and post-fixed in 4% phosphate-buffered paraformaldehyde (PFA) for 6 h before cryoprotection in 30% sucrose in PBS for 24-48 h(Notter et al., 2014; Notter et al., 2018). Brains were cut coronally with a sliding microtome at 30 μm (8 serial sections) and stored at −20 °C in cryoprotectant solution (50 mM sodium phosphate buffer (pH 7.4) containing 15% glucose and 30% ethylene glycol; Sigma-Aldrich, Switzerland) until further processing.

#### Immunofluorescence staining

Immunofluorescence staining was performed according to established protocols(Notter et al., 2014; Notter et al., 2018; Schalbetter et al., 2022). In brief, the sections were washed in Tris-saline buffer and incubated overnight with primary antibodies (**Table 1**) diluted in Tris-saline buffer (0.5 M Tris, 1.5 M NaCl) containing 0.2% Triton-X-100 and 2% normal serum at 4°C under constant agitation (100 rpm). The next day, the sections were washed three times in Tris-saline buffer and incubated for 30 min at room temperature with secondary antibodies (**Table 2**), diluted in Tris-saline buffer containing 2% normal serum, under constant agitation and shielded from light. After incubation, the sections were washed again three times in Tris-saline buffer (shielded from light), mounted on gelatinized glass slides, coverslipped with Dako fluorescence mounting media, and stored at 4°C until image acquisition.

**Table 1:**
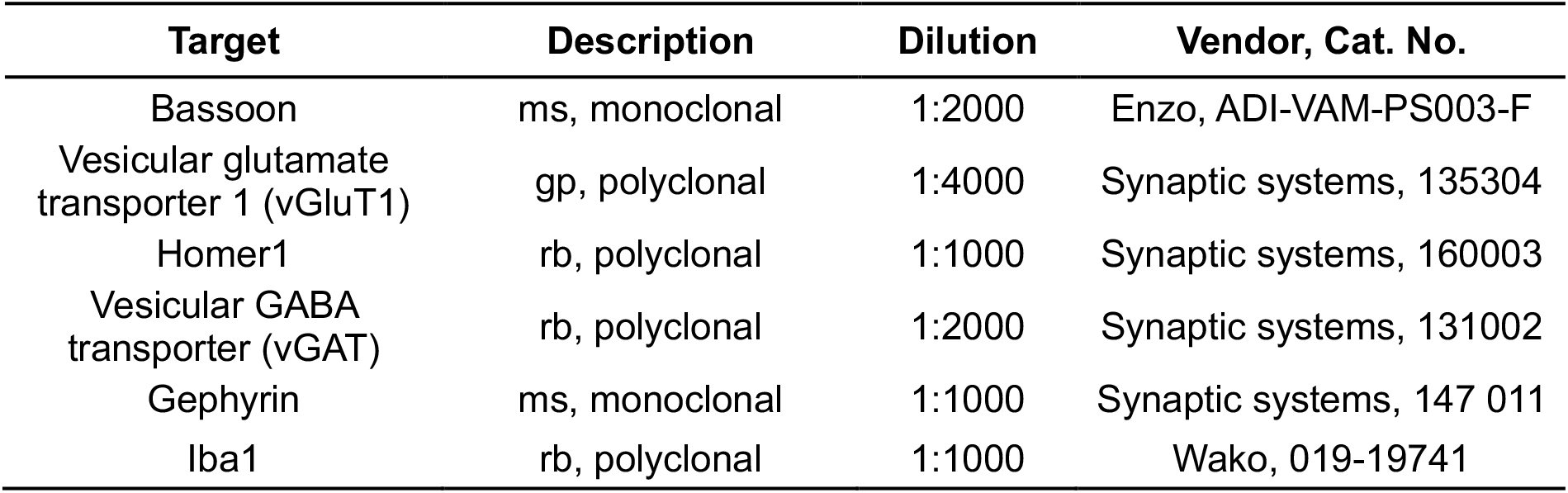
List of primary antibodies used for immunohistochemistry. The selected antibodies have been thoroughly validated in previous studies using C57BL6/N mice(Schalbetter et al., 2022; von Arx et al., 2023). Ms, mouse; gp, guinea pig; rb, rabbit.

**Table 2:**
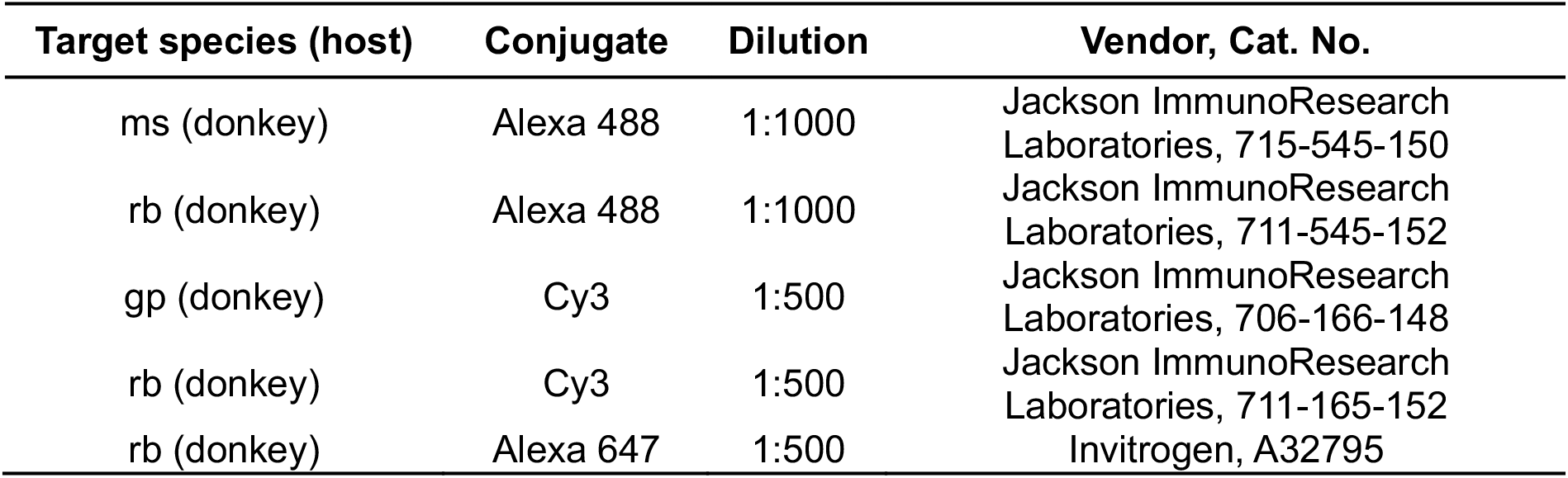
List of secondary antibodies used for immunohistochemistry.

| Target species (host) | Conjugate | Dilution | Vendor, Cat. No. |
| --- | --- | --- | --- |
| ms (donkey) | Alexa 488 | 1:1000 | Jackson ImmunoResearch Laboratories, 715-545-150 |
| rb (donkey) | Alexa 488 | 1:1000 | Jackson ImmunoResearch Laboratories, 711-545-152 |
| gp (donkey) | Cy3 | 1:500 | Jackson ImmunoResearch Laboratories, 706-166-148 |
| rb (donkey) | Cy3 | 1:500 | Jackson ImmunoResearch Laboratories, 711-165-152 |
| rb (donkey) | Alexa 647 | 1:500 | Invitrogen, A32795 |

#### Microscopy and immunofluorescent image analyses

Double immunofluorescence images of vGluT1 and Homer1 (excitatory synapses) or vGAT and Gephyrin (inhibitory synapses) were acquired using a Zeiss LSM900 confocal microscope. Single plane images at a resolution of 2048 × 2048 pixels (pixel size: 0.041 × 0.041 µm) were captured with an oil-immersed 63× objective (NA 1.4, 421782-9900-000, Zeiss). A total of 18 images (6 images per subregion) across three consecutive sections containing the mPFC were captured per animal (*N* = 6 animals per age and sex). All images were collected by an experimenter blinded to the animals’ ages. Due to the data volume, multiple imaging sessions were required. To avoid potential confounds by imaging days, conditions (stained sections from animals of different ages) were randomized across different imaging sessions, so that images from all age groups were acquired on a single imaging day. To estimate the density of excitatory and inhibitory synapses, colocalization between vGluT1^+^ and Homer1^+^ puncta (excitatory synapses) or vGAT^+^ and Gephyrin^+^ puncta (inhibitory synapses) was measured and calculated using a custom-made macro (provided by J.-M. Fritschy, Institute of Pharmacology and Toxicology, University of Zurich, Switzerland) developed for the ImageJ software. This macro has been extensively validated for immunohistochemical colocalization studies and has been described in detail previously(Schalbetter et al., 2022; von Arx et al., 2023). In brief, Gaussian filter, background subtraction, and a threshold were applied to the images for each channel. The settings for each marker were adjusted to achieve an optimal representation of vGluT1, Homer1, vGAT, and Gephyrin, and which were kept constant during image analyses. The number of colocalized clusters was defined as pixel clusters in the presynaptic channel (VGluT1 or VGAT) that overlapped with pixel clusters in the postsynaptic channel (Homer1 or Gephyrin), with a set size cutoff at 0.02 μm^2^. The density (synapses/mm^2^) was calculated by dividing the number of colocalized clusters by the image area (84.51 µm × 84.51 µm). Week-to-week changes in synapse density were assessed by calculating the percent change at each postnatal age relative to the preceding week with the following formula: ((synapse density/mean synapse density of preceding week) × 100)-100.

Immunofluorescence images of sections stained against bassoon were captured using the Olympus IXplore SpinSR10 super resolution imaging system at the Center for Microscopy and Image Analysis (ZMB) of the University of Zurich (UZH). Z-stack images were acquired at a resolution of 2304 × 2304 pixels (pixel size: 0.065 × 0.065 µm, z-step: 0.14 µm) using an oil-immersed 100× objective (NA 1.3, UPLSAPO UPlan S Apo). Nine images within layer II/III of the mPFC (bregma: +2.2 to +1.4 mm) were acquired from three consecutive sections containing the prefrontal subregions anterior cingulate cortex (AC), prelimbic cortex (PrL) and infralimbic cortex (IL) for each animal (with *N* = 10 animals per group and sex except for P21 and P56, where *N* = 6 animals per group and sex. One male in the P28 age group had to be excluded because of bad tissue quality due to ill perfusion). The density of Bassoon-positive presynaptic boutons was estimated using the ImageJ software. For this purpose, an automated threshold (“moments dark”) and the watershed function were applied to z-projected images. Bassoon puncta were then counted with the particle count function. Density (puncta/mm^2^) was calculated by dividing the number of particles with the area of the image (149.76 µm × 149.76 µm) × 1’000’000. Week-to-week changes in synapse density were assessed by calculating the percent change at each postnatal age relative to the preceding week with the following formula: ((density (puncta/mm^2^)/mean density of preceding week) × 100)-100.

Double-immunofluorescence images of sections stained against the microglia marker Iba1 and Bassoon were captured using the Olympus IXplore SpinSR10 super-resolution imaging system (ZMB, UZH). An oil-immersed 100× objective (NA 1.3, UPLSAPO UPlan S Apo) was used. Images were acquired at a resolution of 2304 × 2304 pixels (pixel size: 0.065 × 0.065 µm, z-step: 0.27 µm). Three images per region (AC, PrL, IL) were obtained, resulting in a total of 9 images across the mPFC from three consecutive sections per animal (with *N* = 6 animals per age group and sex. One male in the P28 age group had to be excluded because of bad tissue quality due to ill perfusion). All images were deconvoluted with Huygens Professional version 23.04 using the CMLE algorithm with SNR:10 and 40 iterations (Scientific Volume Imaging, The Netherlands). After deconvolution, the images were imported into Imaris image analysis software (version 10.2.0, Oxford Instruments). Iba1^+^ microglia were reconstructed three-dimensionally using the “surface” creation. Bassoon^+^ synaptic puncta were generated using the “spots” creation. The “Find spots close to surface” with a threshold of 0 (determining whether the center of the Bassoon^+^ spots is inside or outside the microglia surface) was used to quantify Bassoon^+^ synaptic particles within surface-rendered Iba1^+^ microglia. The phagocytic index was calculated by dividing the number of synaptic particles with the volume of microglia within each image. An average of 47 ± 4 (mean ±SD, males) and 45 ± 5 (mean ±SD, females) microglia were analyzed per age group. The average volume of microglia was estimated by dividing the total volume of microglia by the number of microglia analyzed per image. The density of microglia (number/mm^2^) was calculated by dividing the number of microglia with the area of the image (149.76 µm × 149.76 µm) × 1’000’000.

### Blinding and statistical analyses

All data were collected and analyzed blindly, except for the spine analysis, where the previous imaging day needed to be confirmed to accurately quantify spine dynamics. All statistical analyses were performed using Prism (version 10.6.1, GraphPad Software, La Jolla, CA, USA). Statistical significance was set at *p* < 0.05. All data met the assumptions of normal distribution and equality of variance. Because the dataset of the longitudinal study contained missing values due to the described dropouts, a mixed-effects model using residual maximum likelihood (REML) estimation was used. Synaptic density across different ages in the cross-sectional experiments was assessed using one-way ANOVA or two-way ANOVA. Percent change values derived from the week-to-week synapse density comparisons were evaluated using one-sample *t*-tests, with 0 serving as the baseline representing no change relative to the previous week. For all analyses, females and males were analyzed separately, given previously reported sex-specific differences in synaptic remodeling in the rat mPFC(Drzewiecki et al., 2016). Whenever appropriate, ANOVAs or mixed-effects models were followed by Tukey’s or Šídák’s post-hoc test for multiple comparisons.

## ACKNOWLEDGMENTS

This work was financially supported by the Swiss National Science Foundation (grant No. PZ00P3_202149 awarded to T.N.).

## CONTRIBUTIONS

J.F., S.F., A.v.F.C., S.M.S, V.B., E.B., and M.T.W. were involved in the acquisition, analysis, and interpretation of the study data; T.N. was involved in the conception and design of the study and analysis and interpretation of the study data; T.N., B.W., and U.M. supervised research; T.N. and U.M. wrote the initial manuscript draft; all authors contributed to the reviewing and editing of the manuscript, and have given final approval for the version to be published.

## DATA AVAILABILITY

All data are available in the main text or the supplementary information. Source data are provided with this paper.

