## Supplementary information for "Developmental Trajectory of Synaptic Remodeling in the Mouse Prefrontal Cortex"

### SUPPLEMENTARY DATA

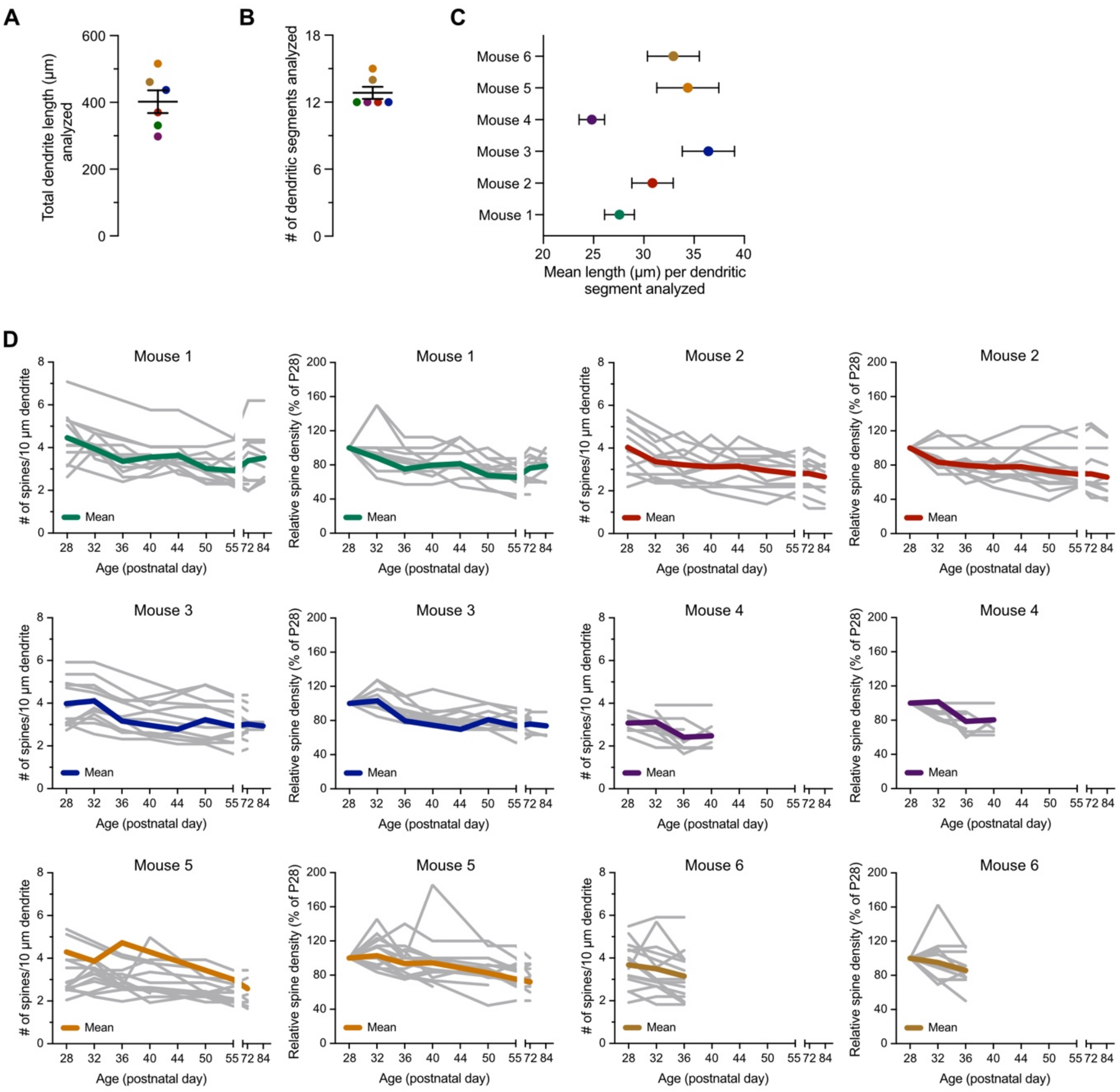


**Supplementary Figure S1. Longitudinal high-resolution *in vivo* two-photon imaging. (A)** Total dendritic length analyzed per animal (each dot represents a different animal). **(B)** Number of dendritic segments analyzed per animal (each dot represents a different animal). **(C)** Mean length of dendritic segment analyzed per animal (each dot represents a different animal). **(D)** Number of spines per 10 µm dendrite (left) and relative spine density (% of P28, right) across the development of the mPFC for each mouse. Colored lines represent the means of spine changes, while individual grey lines represent measurements from each analyzed dendritic segment. Longitudinal imaging in three mice (mice 4, 5, and 6) could not be completed to the final age (P84) due to a shift in the microprism.

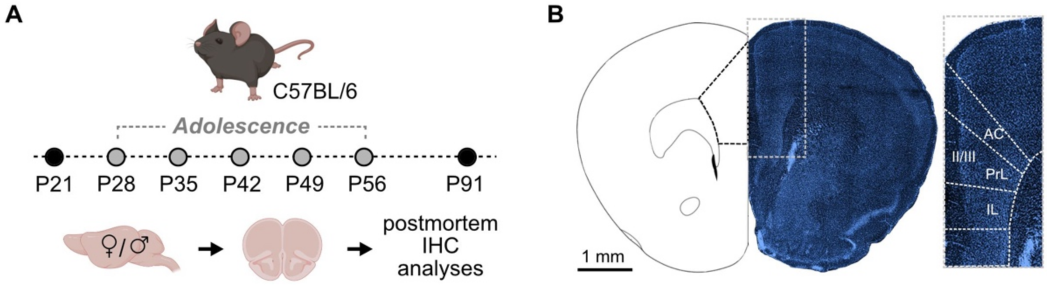


**Supplementary Figure S2. Cross-sectional analyses of synaptic densities across postnatal maturation in C57BL6/N mice. (A)** Cross-sectional immunohistochemical analyses were performed on prefrontal tissue collected from C57BL/6N male and female mice at postnatal days (P) 21, 28, 35, 42, 49, 56, and 91, spanning juvenile (preadolescent, black dot), adolescent (grey dots), and adult developmental stages (black dot). **(B)** Representative tile scan image of a brain section containing mPFC subregions (anterior cingulate (AC), prelimbic (PrL), infralimbic (IL)) stained with DAPI. All postmortem evaluations were conducted on images acquired from layer II/III spanning all three prefrontal subregions.

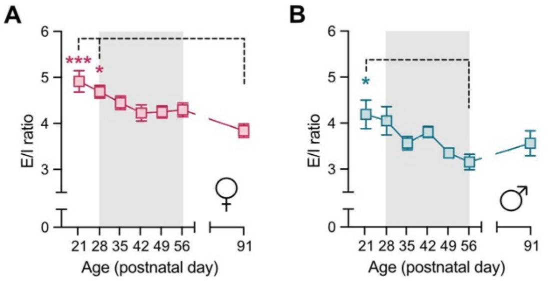


**Supplementary Figure S3. Developmental trajectory of excitatory/inhibitory (E/I) balance across prefrontal maturation. (A)** E/I ratio across postnatal maturation in female C57BL6/N mice. ***$p$ < 0.001, *$p$ < 0.05 based on post-hoc test following one-way ANOVA. **(B)** E/I ratio across postnatal maturation in male C57BL6/N mice. *$p$ < 0.05 based on post-hoc test following one-way ANOVA. All data are means ± SEM. $N$ = 6 mice per group and age.

**Cross-sectional analyses of synaptic density using immunohistochemistry**

Sample collection, immunofluorescence staining and image acquisition were performed according to established protocols [1,7,8]. Tissue was collected from C57BL/6N male and female mice at P21, P28, P35, P42, P49, P56, and P91, spanning preadolescent, adolescent, and adult developmental stages [9] (**Supplementary Fig. 2**). These time points were selected to achieve high temporal resolution of synaptic remodeling across adolescence [10] and to capture potential sex-dependent differences related to pubertal onset [10,11], which occurs between P30–P32 in females and P35–P40 in males [12,13].

*Sample collection:* Animals were deeply anesthetized with an overdose of pentobarbital (Esconarkon ad us. vet., Streuli Pharma AG, Switzerland) and perfused transcardially with artificial cerebrospinal fluid (aCSF, pH 7.4) with a perfusion rate of 20 ml/min. The brains were immediately removed from the skull and post-fixed in 4% phosphate-buffered paraformaldehyde (PFA) for 6 h before cryoprotection in 30% sucrose in PBS for 24-48 h[1,8]. Brains were cut coronally with a sliding microtome at 30 µm (8 serial sections) and stored at −20 °C in cryoprotectant solution (50 mM sodium phosphate buffer (pH 7.4) containing 15% glucose and 30% ethylene glycol; Sigma-Aldrich, Switzerland) until further processing.

*Immunofluorescence staining:* Immunofluorescence staining was performed according to established protocols [1,7,8]. In brief, the sections were washed in Tris-saline buffer and incubated overnight with primary antibodies (**Supplementary Table 1**) diluted in Tris-saline buffer (0.5 M Tris, 1.5 M NaCl) containing 0.2% Triton-X-100 and 2% normal serum at 4°C under constant agitation (100 rpm). The next day, the sections were washed three times in Tris-saline buffer and incubated for 30 min at room temperature with secondary antibodies (**Supplementary Table 2**), diluted in Tris-saline buffer containing 2% normal serum, under constant agitation and shielded from light. After incubation, the sections were washed again three times in Tris-saline buffer (shielded from light), mounted on gelatinized glass slides, coverslipped with Dako fluorescence mounting media, and stored at 4°C until image acquisition.

**Supplementary Table 1: List of primary antibodies used for immunohistochemistry.** The selected antibodies have been thoroughly validated in previous studies using C57BL6/N mice [7,14]. Ms, mouse; gp, guinea pig; rb, rabbit.

| Target | Description | Dilution | Vendor, Cat. No. |
|---|---|---|---|
| Bassoon | ms, monoclonal | 1:2000 | Enzo, ADI-VAM-PS003-F |
| Vesicular glutamate transporter 1 (vGluT1) | gp, polyclonal | 1:4000 | Synaptic systems, 135304 |
| Homer1 | rb, polyclonal | 1:1000 | Synaptic systems, 160003 |
| Vesicular GABA transporter (vGAT) | rb, polyclonal | 1:2000 | Synaptic systems, 131002 |
| Gephyrin | ms, monoclonal | 1:1000 | Synaptic systems, 147 011 |
| Iba1 | rb, polyclonal | 1:1000 | Wako, 019-19741 |

**Supplementary Table 2: List of secondary antibodies used for immunohistochemistry.**

| Target species (host) | Conjugate | Dilution | Vendor, Cat. No. |
|---|---|---|---|
| ms (donkey) | Alexa Fluor 488 | 1:1000 | Jackson ImmunoResearch Laboratories, 715-545-150 |
| rb (donkey) | Alexa Fluor 488 | 1:1000 | Jackson ImmunoResearch Laboratories, 711-545-152 |
| gp (donkey) | Cy3 | 1:500 | Jackson ImmunoResearch Laboratories, 706-166-148 |
| rb (donkey) | Cy3 | 1:500 | Jackson ImmunoResearch Laboratories, 711-165-152 |
| rb (donkey) | Alexa Fluor 647 | 1:500 | Invitrogen, A32795 |
